# Nucleoporin NUP210L and BAF-paralogue BAF-L together ensure microtubule organization and nuclear integrity in spermatids

**DOI:** 10.1101/2024.05.16.594580

**Authors:** Maha Al Dala Ali, Guy Longepied, Nicolas Lévy, Catherine Metzler-Guillemain, Michael J Mitchell

**Author notes:** Correspondence should be addressed to M J Mitchell;. Present address: Servier, Research and Development, rue Francis Perrin, 91190 Gif-sur-Yvette (Nicolas Lévy).

## Abstract

During spermiogenesis, haploid round spermatids differentiate into spermatozoa. This involves nuclear elongation, chromatin compaction, cytoplasm reduction, and formation of the acrosome and the flagellum. These events are orchestrated by cytoskeletal elements - acroplaxome and manchette - that attach to the nuclear envelope (NE) except at the caudal pole where the nuclear pore complexes (NPCs) shift to form a dense array. Here, we use a genetic dissection approach to reveal function at the caudal NE, through the study of two spermatid-specific proteins, whose human orthologues persist there as spermatids elongate: NUP210L, a transmembrane nucleoporin and BAF-L, paralogue and interactor of chromatin protein BAF. In mice, inactivation of either BAF-L or NUP210L has no impact on fertility. However, we show here that in mice lacking NUP210L, two copies of BAF-L become essential for fertility; in *Nup210l*^−/−^,*Banf2*^+/−^ or *Nup210l^−/−^*,*Banf2*^−/−^ mice, most spermatids arrest during nuclear elongation (step 10-11) with mislocalized NPCs and disorganized manchette microtubules that frequently invaginate the nucleus from the caudal pole. Our results suggest that the NPC array, and BAF-L/BAF, ensure nuclear integrity at the caudal pole during spermatid remodeling.

**Summary:** Nucleoporin NUP210L and BAF-paralogue BAF-L function redundantly during mouse spermatid nuclear remodeling to concentrate nuclear pores to the flagellar pole, organize the manchette cytoskeleton and prevent nuclear invagination by microtubules.

## Introduction

Spermiogenesis is the post-meiotic phase of spermatogenesis when haploid round spermatids transform into specialized motile spermatozoa able to transport the paternal genetic information into the oocyte cytoplasm and create the zygote. To accomplish this, spermatids develop a flagellum for motility and an acrosome to penetrate the oocyte’s protective *zona pellucida* layer. To aid motility, and protect the DNA from damage, the nucleus is elongated and condensed. Nuclear compaction involves re-packaging of the chromatin into a dense inactive form by the replacement of almost all histones with small basic proteins, the protamines.

This unique nuclear metamorphosis is structured by cytoskeletal scaffolding that all but surrounds the nucleus (Kierszenbaum and Tres, 2004): the acroplaxome under the acrosome, which is bounded by the perinuclear ring from which descends the manchette, a sleeve of microtubules attached to the lateral nuclear membranes by links involving heterodimers of SUN3 and SUN4 inner nuclear membrane proteins (Calvi *et al*., 2015; Gao *et al*., 2020; Pasch *et al*., 2015). In the mouse, the manchette forms in round spermatids at step 8, just prior to the beginning of elongation, and disappears after both elongation and condensation of the nucleus have occurred at step 13-14 (Lehti and Sironen, 2016). The acroplaxome-acrosome attachment to the nucleus is dependent on the inner nuclear membrane protein DPY19L2 (Pierre *et al*., 2012). In mice lacking SUN3, SUN4 or DPY19L2 the manchette does not attach to the nucleus and all spermatozoa have compacted round heads, showing that nuclear attachment of the acroplaxome and the manchette are required for nuclear elongation and shaping but not compaction (Calvi *et al*., 2015; Gao *et al*., 2020; Pasch *et al*., 2015; Pierre *et al*., 2012).

In mouse and human, major changes occur to the NE during spermatid elongation with the dismantling and removal of the nuclear lamina ahead of the spreading acroplaxome (Alsheimer *et al*., 1998; Schütz *et al*., 2005; Elkhatib *et al*., 2015; Ho 2010). When the manchette forms, NPCs move from the lateral NE to the caudal pole of the nucleus and form a tight array there, but are excluded from the basal plate where the centrioles and the flagellum attach to the nucleus (Dooher and Bennett 1973; Ho 2010). How NPCs shift from under the manchette is unknown, but it could involve SUN1 which moves to the caudal pole at the same time (Göb *et al*., 2010). *Sun1*-knockout male mice produce no spermatids due to a meiotic block, and so how SUN1 contributes to spermiogenesis has not yet been investigated (Chi *et al*., 2009; Ding *et al*., 2007). Since the NPC is the major non-vesicular route in and out of the nucleus, the caudal pole must be a site of intense trafficking across the nuclear membrane during nuclear remodelling. Nevertheless, in contrast to the rest of the spermatid NE, little else is known about organisation and function within the zones of cytoplasm, NE or nucleoplasm that juxtapose the nuclear attachment site of the flagellum.

Evidence that the NPC plays an important role in spermatid nuclear compaction and remodelling has recently emerged from a genetic study of human male infertility. A biallelic loss-of-function (LoF) mutation in the gene encoding NUP210L, a spermatid-predominant paralogue of the transmembrane nucleoporin NUP210, was identified in an infertile man with a unique phenotype characterized by spermatozoa with a large uncompacted head and histonylated chromatin (Arafah *et al*., 2021). We have shown that NUP210L localizes to the caudal nuclear pole of elongating spermatids and spermatozoa, in mouse and human (Al Dala Ali *et al*., 2024). Its paralogue NUP210 is thought to form an octamer ring around the nuclear pore membrane, that may optimize nuclear pore channel function by buffering it from external forces within the NE (Zhang *et al*., 2020). The inactivation of *Nup210l* alone in the mouse does not, however, impact fertility or chromatin condensation, although spermatozoa do show morphological changes in the head and flagella, and reduced numbers are motile (Al Dala Ali *et al*., 2024). Thus, whether NUP210L is essential for nuclear remodeling and chromatin condensation during human spermiogenesis remains an open question.

We have previously shown that, in human spermatids, the small conserved chromatin protein BAF, and its spermatid-predominant paralogue BAF-L, shift from the nucleoplasm to the caudal nuclear pole as nuclear elongation begins, suggesting that they contribute to nuclear remodelling (Elkhatib *et al*., 2017). In somatic cells, BAF exists as homodimers that when dephosphorylated can bind non-specifically to double stranded DNA, promoting bridging and compaction of DNA (Skoko *et al*., 2009; Zheng *et al*., 2000). In somatic cells, BAF-DNA interactions are at the root of fundamental cellular processes such as nuclear membrane reassembly at the end of mitosis, NE repair, gene regulation, the innate immune response to cytosolic DNA and the DNA damage response (Sears and Roux, 2020). In contrast to BAF, little is known about BAF-L function except that it is unable to bind DNA, can form heterodimers with BAF and has reduced affinity for the LEM-domains bound by BAF (Tifft *et al*., 2006). It has been proposed that BAF-L may regulate BAF function in spermatids but three independent studies, plus the present study, find that males lacking BAF-L are fertile, (Niu *et al*., 2021; Huang *et al*., 2022; Lu *et al*., 2019) providing no insights into the role of BAF-L. Here we present mice carrying null alleles for both BAF-L and NUP210L. We created these mice to look for functional interactions between BAF-L and NUP210L, based on their co-localisation at the caudal pole in human elongating spermatids (Elkhatib *et al*., 2017; Al Dala Ali *et al*., 2024). In stark contrast to the fertility of the single knockouts of BAF-L or NUP210L, we find that male mice lacking NUP210L are sensitive to haploinsufficiency for BAF-L; males are infertile, have a striking deregulation of microtubule organisation during nuclear elongation and produce only low numbers of immotile spermatozoa, most with abnormal but compacted heads. Our study shows that, in spermatids, BAF-L and NUP210L are mutually redundant components of an important nexus between the chromatin, the NPC and the cytoskeleton, bringing function at the caudal nuclear pole of the elongating spermatid into sharp focus.

## Results

### Normal fertility and testicular histology in mice lacking BAF-L

To investigate the biological function of BAF-L in vivo, we used CRISPR-Cas9 technology to target its coding gene, *Banf2,* and created a mouse line carrying the null allele, *Banf2^em42Mmjm^,* hereafter referred to as *Banf2*^−^. The *Banf2*^−^ allele has a 23 base pair (bp) deletion in exon 3 that includes the initiation codon (based on transcript NM_1178477.5 - Fig. S1A). Purification and sequencing of RT-PCR products from wild type and *Banf2^−/−^* adult testis revealed that the 23 bp deletion induces the skipping of exon 3 that encodes the N-terminal half of BAF-L (42 of 90 amino acids) (Fig. S1B). In exon 4, there is one in-frame ATG at codon 53 but it is in a weak Kozak context, ctgATGc, indicating that no part of BAF-L is expressed from our *Banf2^−^* allele. As in previously published reports of *Banf2*-KO mice (Huang *et al*., 2022; Lu *et al*., 2019; Niu *et al*., 2021), our *Banf2^−/−^* male mice are fertile. We found them to have normal litter frequency and litter size (Fig. S2A). Testis size and weight were normal (Fig. S2C) No histological anomalies of spermatogenesis were evident (Fig. S2D), and spermatozoa had normal morphology, count and motility (Fig. S2E, F and G).

### Two copies of *Banf2* are required for male fertility in mice lacking *Nup210l*

Nuclear pore complexes reorganize into a tight array at the caudal nuclear pole, as the manchette forms and spermatid elongation initiates, and we have shown that this is also the case for BAF, BAF-L, the spermatid-specific nucleoporin NUP210L and the widely expressed nucleoporin NUP153 (Al Dala Ali *et al*., 2024; Elkhatib *et al*., 2017), indicating functional coordination, although mice lacking either BAF-L or NUP210L are fertile. As part of an effort to identify function at the spermatid posterior pole, we generated mice carrying null alleles for both *Banf2* and *Nup210l*.

We first tested the fertility of mice carrying three combinations of null alleles for *Nup210l* and *Banf2* (*Banf2*^−/−^, *Nup210l*^+/−^; *Banf2*^+/−^,*Nup210l*^−/−^; *Banf2*^−/−^,*Nup210l*^−/−^). Five males of each genotype and 3 double heterozygote male controls (*Banf2*^+/−^;*Nup210l*^+/−^) were paired with two WT females over a period of two months. The average litter size for each genotype is shown in Table 1.

**Table 1:** Fertility test of *Banf2,Nup210l* mutant males.

| Genotype | | Fertile males/Total males | Litter size (mean ± SEM) |
| --- | --- | --- | --- |
| *Banf2* | *Nup210l* |  |  |
| +/- | +/- | 3/3 | 12.0 ± 1 |
| -/- | +/- | 5/5 | 11.7 ± 0.6 |
| +/- | -/- | 0/5 | 0 |
| -/- | -/- | 0/5 | 0 |

Mice with zero copies of wild-type *Nup210l* and one or zero copies of wild-type *Banf2* were infertile (Table 1). Thus, in mice lacking NUP210L function, male fertility is sensitive to reduced expression of BAF-L, while in mice with reduced expression of BAF-L, NUP210L function is critical for fertility.

### Lack of BAF-L and NUP210L affect sperm morphology and concentration

We next investigated the nature and the cause of the infertility in *Banf2*^+/−^,*Nup210l*^−/−^ and *Banf2*^−/−^,*Nup210l*^−/−^ males, through the analysis of testicular and epididymal histology, and epididymal spermatozoa. Testis weight in adult infertile mutants (2 months old) was not significantly different to that of the control group (Fig. 1A, 1C). However, spermatogenic activity was clearly disrupted in infertile double mutant males, as evidenced by abnormally low numbers or complete absence of spermatozoa in the epididymis (Fig. 1B).

**Figure 1:**
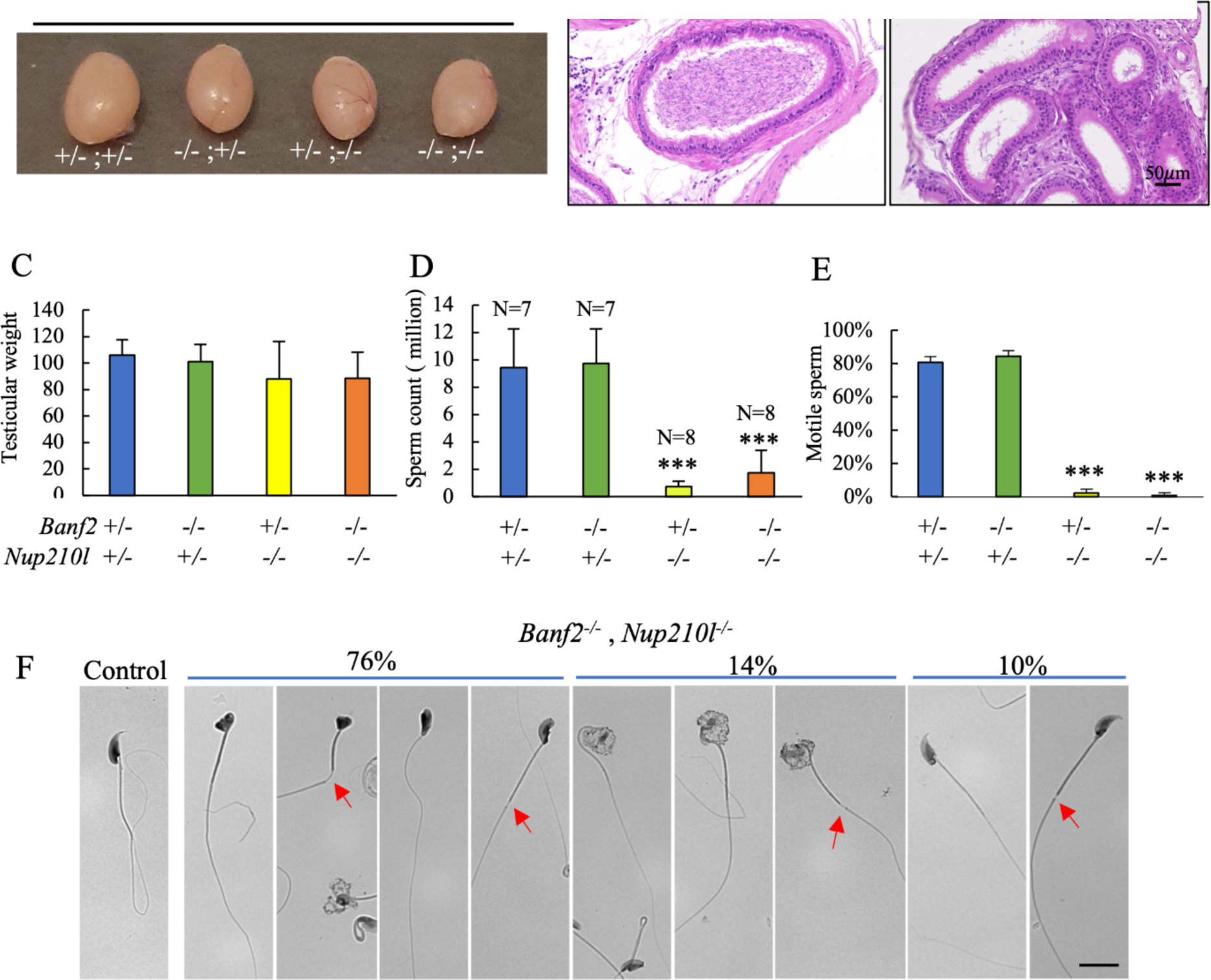
Spermatogenesis disruption in *Banf2,Nup210l* mutant mice. (A) Representative images of adult testes from the indicated genotypes (B) Sections of epididymis stained with hematoxylin and eosin (H&E) from control and *Banf2^−/−^,Nup210l^−/−^* mice (C) The average weight of adult testes in *Banf2^+/−^,Nup210l*^−/−^ and *Banf2^−/−^,Nup210l^−/−^* males (n=8) is comparable to the control group (n=7); *p*-value ≥ 0,08 (D) Sperm count was significantly reduced in both *Banf2^+/−^,Nup210l^−/−^* and *Banf2^−/−^,Nup210l^−/−^* while *Banf2^−/−^,Nup210l^+/−^* have normal sperm count. *p*-value < 0.00001, 0.00001, and 0,82 respectively. (E) Quantification of the number of motile sperm shows that the majority of spermatozoa produced by *Banf2^−/−^; Nup210^l−/−^* and *Banf2^−/−^;Nup210l^−/−^* were immotile *(p*-value <0.00001, 0.00001). (F) Morphological abnormalities of epididymal sperm in both *Banf2^−/−^,Nup210^l−/−^* and *Banf2^−/−^,Nup210l^−/−^* range from abnormal head shape (76%), to a large decondensed sperm head (14%), and only 10% have the usual falciform morphology. Scale bar 10 μm.

To assess the quality of sperm production, epididymal sperm samples were collected from 16 double mutant males of the *Banf2*^+/−^,*Nup210l*^−/−^ or *Banf2*^−/−^,*Nup210l*^−/−^ genotype. Compared to control littermates (n=7) the sperm count was significantly lower in the double mutants (Fig 1D): five had less than 30%, nine less than 10%, and two had no spermatozoa. Furthermore, 98% of the sperm in the mutant mice were immotile (Fig. 1E). Morphological examination revealed that 90% of sperm have an abnormal head shape, with 76% losing their hook-like characteristic shape, and 14% exhibiting a large uncompacted head (Fig. 1F). Additionally, we observed tail abnormalities characterized by midpiece defects (Fig. 1F). These findings show that reduced expression of BAF-L in the absence of NUP210L impacts spermatozoa production and morphogenesis.

### Spermatids arrest during elongation in *Nup210l*^−/−^ mice carrying a *Banf2* null allele

Histological analysis of testes stained with hematoxylin and Periodic Acid-Schiff (H-PAS) showed that meiosis and spermatid development were normal up to step 10 of spermiogenesis. In contrast, the number of elongated spermatids at later steps was greatly reduced in *Banf2*^+/−^,*Nup210l*^−/−^ and *Banf2*^−/−^,*Nup210l*^−/−^ males (Fig.S2). This was most evident in stage I-VIII tubules, where condensed elongated spermatids are normally abundant (Fig. 2A). Using the TUNEL assay, in three biological triplicates, we found a 30-fold increase in apoptotic cells in the seminiferous tubules of *Banf2*^−/−^,*Nup210l*^−/−^ versus control males (185 ± 12 per 20 tubules versus 6 ± 1 per 20 tubules, Fig. 2B, 2C). Most apoptotic cells were uncondensed elongated spermatids, identified by both their morphology and their location within seminiferous tubules (Fig. 2B). Our results suggest that, in *Banf2*^+/−^,*Nup210l*^−/−^ and *Banf2*^−/−^,*Nup210l*^−/−^ males, the majority of spermatids fail to progress beyond early elongation steps, show no signs of nuclear condensation and are eliminated by apoptosis prior to step 13.

**Figure 2:**
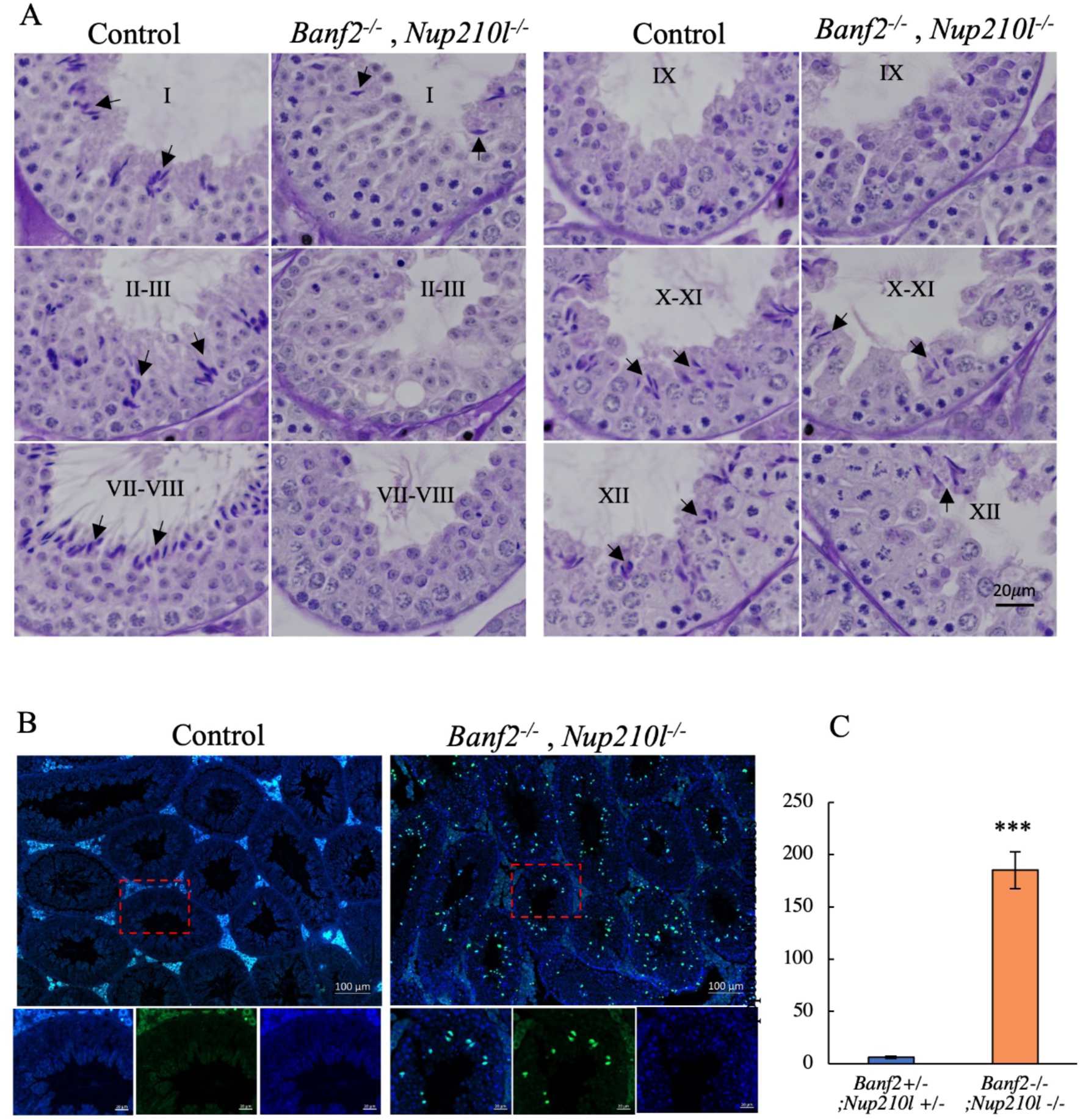
Significant loss of elongated spermatids in *Banf2^−/−^,Nup210l^−/−^* mice. (A) staged comparison of H-PAS-stained *Banf2^−/−^,Nup210l^−/−^* and control littermate testes; Roman numbers represent the stages of the seminiferous tubules. Black arrows mark the elongated spermatid. A noticeable decrease in the number of elongated spermatids is observed in *Banf2^−/−^,Nup210l^−/−^* tubules. (B) Images of TUNEL-stained mouse testis sections, with green indicating TUNEL-positive cells. The number of apoptotic cells has significantly increased in mutant mice. Upon closer examination (magnification of the red boxed area), it is evident that the majority of apoptotic cells are elongated spermatid cells. (C) Quantifications of TUNEL-labeled cells revealed a significant increase in apoptosis in mutant testes. *p*-value = 0.00007.

### Inefficient chromatin remodeling in infertile double mutant mice

To investigate whether histone-to-protamine replacement is affected in *Banf2^+/−^,Nup210l^−/−^* and *Banf2^−/−^,Nup210l^−/−^* males, we initially determined chromomycin A3 (CMA3) staining of epididymal spermatozoa. CMA3 is a fluorochrome specific for guanine-cytosine (GC)-rich sequences in histonylated chromatin, but its binding is blocked when the chromatin is packaged with protamines. Normal spermatozoa exhibit a negative CMA3 test result, and a positive result indicates histone retention and protamine deficiency. In both *Banf2*^+/−^,*Nup210l*^−/−^ and *Banf2*^−/−^,*Nup210l*^−/−^ males, we observed that approximately 22-25% of the sperm were CMA3 positive compared to only 1% in controls (Fig. 3 A and B).

**Figure 3:**
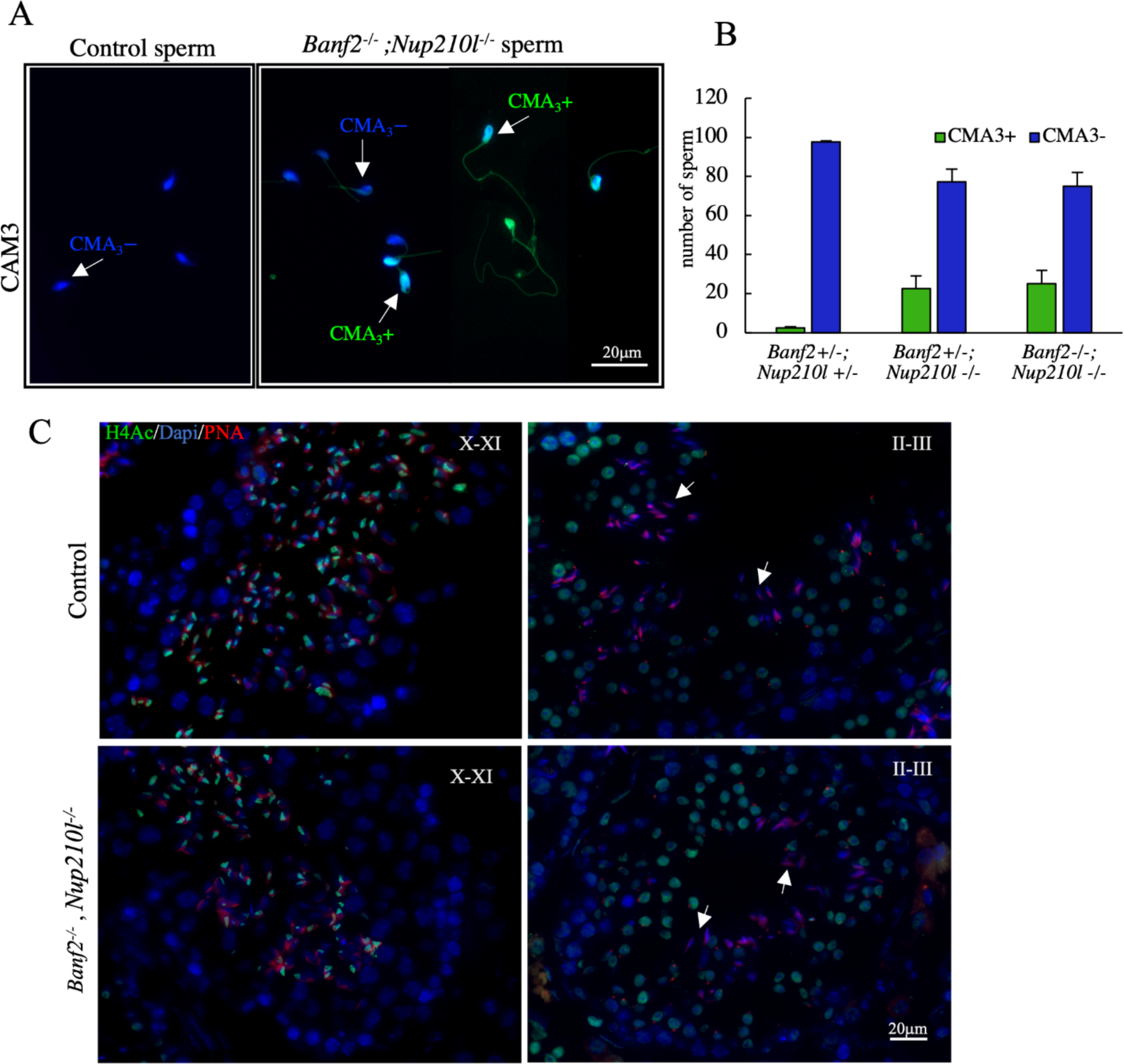
Inefficient histone eviction during spermiogenesis in the *Banf2^−/−^,Nup210l^−/−^*. (A) Chromomycin A_3_ (CMA_3_) was used to stain mouse sperm; CMA_3_+ indicates positive spermatozoa (bright green); CMA_3_-indicates negative spermatozoa. (B) Quantification of the CMA_3_+ and CMA_3_-spermatozoa reveals a significant increase in CMA_3_+ sperm in the *Banf2^−/−^,Nup210l^−/−^*, with a *p*-value= 0.005. (C) Immunoflourescence with antibody specific to acetylated Histone 4 (H4Ac) is performed on testis sections from mutant and control littermates.

We then assessed histone H4 acetylation by immunofluorescence. The timing and progression of H4 acetylation in the mutant animals appeared normal, beginning at spermatid step 8 and intensifying at step 9 and 10, before receding to the caudal pole at step 11 and 12 and disappearing in later stages (Fig.3 C). Although the pattern of acetylated histone 4 signal appears normal up to the block, the presence of significant number of CMA3 positive cells in the epididymis of double mutants indicates that the histone-to-protamine replacement is perturbed in some but not all spermatids.

### Loss of NE integrity in round spermatids of *Banf2^−/−^,Nup210l^−/−^* males

To investigate the form of the NE and the evolution of the nuclear lamina (NL) in round spermatids in *Banf2*^−/−^,*Nup210l*^−/−^ males, we used an antibody against lamin B1. In controls and double-KO mice, we observed the normal pattern of labelling (Pierre *et al*., 2012), with lamin B1 at the nuclear periphery, except under the developing acrosome in round spermatids until its disappearance at the initiation of nuclear elongation, indicating that the absence of BAF-L and NUP210L does not affect the dismantling of the NL in round spermatids. Nevertheless, these analyses revealed a major difference in the shape of the round spermatid nucleus: in controls it was round, but in *Banf2*^−/−^,*Nup210l*^−/−^ males, beginning at step 1, it was deformed, appearing elliptical, angular or indented (Fig. 4A). No differences were obvious later in step 9-10 spermatids, with both control and *Banf2*^−/−^,*Nup210l*^−/−^ showing uniform elongated nuclei. These observations indicate that BAF-L and NUP210L are required to maintain a spherical NE in round spermatids. Interestingly, we did not observe deformity of the NE in disaggregated spermatids (Fig. S4), implying that, in the testis, round spermatid nuclei lacking BAF-L and NUP210L are being deformed by extracellular forces exerted from within the seminiferous epithelium.

**Figure 4:**
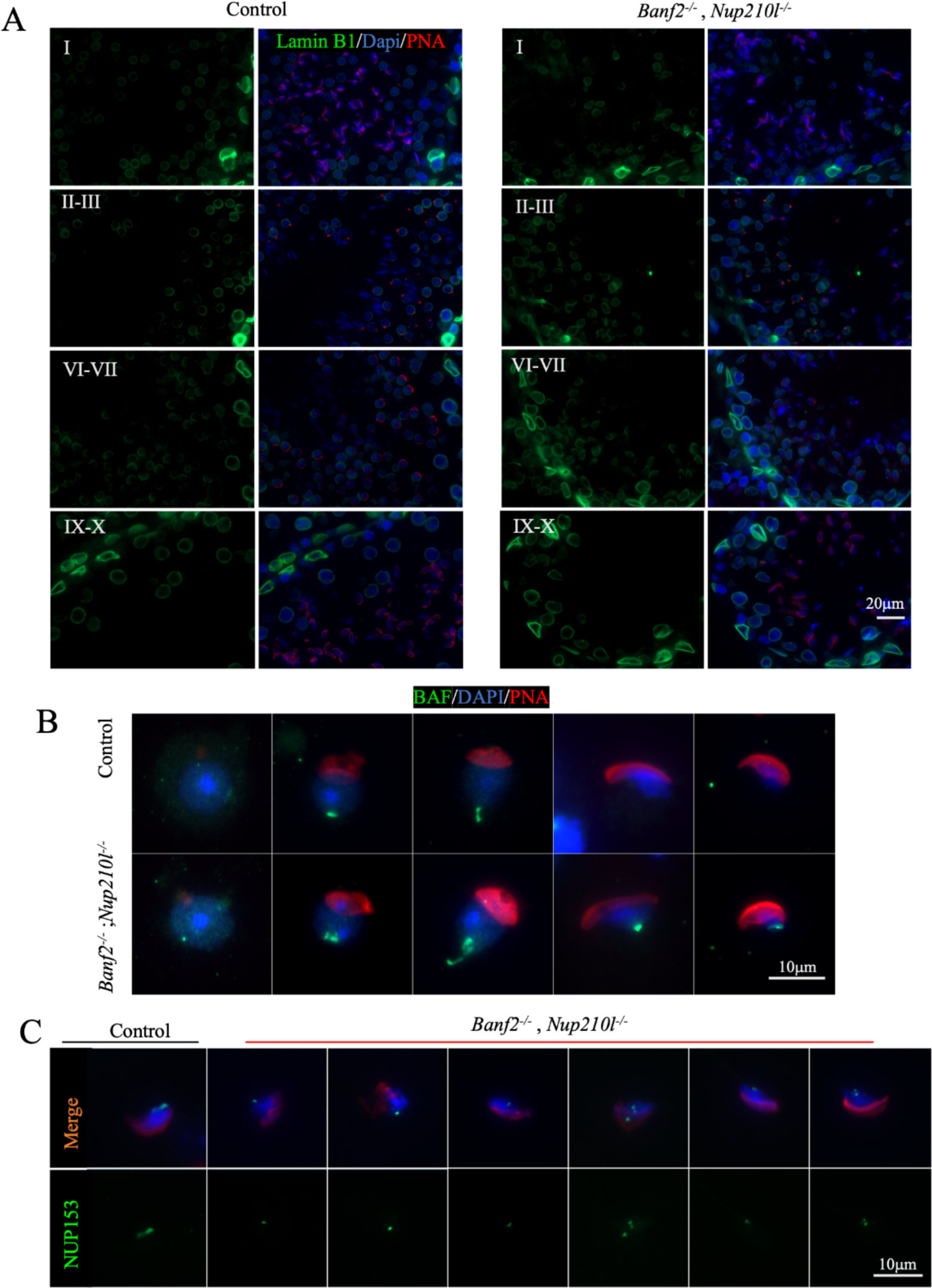
*Banf2^−/−^,Nup210l^−/−^* spermatids exhibit disorganization of NE components. (A) immunofluorescence of testis sections from mutant and control littermates with antibody against Lamin B1 (green), nuclei counterstained with DAPI (blue) and acrosome by PNA (red) as an acrosomal marker to identify the stages of spermatogenesis*. (*B) Immunofluorescence assessment of BAF localization using a polyclonal antibody against BAF (green), nuclei labeled with DAPI (blue), and acrosome by PNA (red) (C) nuclear pore complex (NPC) localization was detected with a specific antibody against NUP153 (green); nuclei were counterstained with DAPI and acrosomes with PNA.

### BAF shows increased nuclear retention in *Banf2*^−/−^,*Nup210l*^−/−^ round spermatids

Since BAF-L can heterodimerize with BAF *in vitro*, we investigated how the absence of BAF-L affected BAF localisation in *Banf2*^−/−^,*Nup210l*^−/−^ spermatids by immunofluorescence. In control mice, BAF exhibits cytoplasmic and nucleoplasmic localization in step 1-3 round spermatids. BAF becomes exclusively nucleoplasmic in spermatids from step 4-5, concentrated at a point on the nuclear periphery at the posterior pole, where the distal centriole and the flagellar axoneme contact the nucleus. In step 9 elongated spermatids, it is only present at the caudal pole next to the flagellum attachment site. BAF is not detected in spermatids after step 9 (Fig. 4B). In epididymal sperm, there is a complete absence of BAF signal (Fig. S4).

In *Banf2^−/−^,Nup210l^−/−^* males, unlike controls, BAF predominates in the nucleus in step1-3 round spermatids (Fig. 4B). The specific localisation of BAF at the caudal pole in step 9 spermatids is as in controls, but unlike in controls, BAF persists at the caudal pole in later elongating spermatids (Fig. 4B) and is still present in 23% of epididymal spermatozoa (n=100) (Fig. S4).

### Nuclear pore array disorganised at the caudal pole in *Banf2*^−/−^,*Nup210l*^−/−^ spermatids

NUP210L is thought to be a component of the NPC in spermatids. We therefore next investigated the localization of the NPC in *Banf2*^−/−^,*Nup210l*^−/−^ spermatids using an antibody against NUP153, a component of the NPC present in spermatids. Normally, the NPC is located at the posterior nuclear pole surrounding the basal plate where the flagellum connects to the nucleus. In control spermatids, NUP153 is detected along the caudal limit of the nucleus distal to the acrosome. However, in the *Banf2*^−/−^,*Nup210l*^−/−^ males, we observed altered NPC localization in elongating spermatids, where the NUP153 signal was either concentrated at the ventral part of the caudal pole or completely displaced from the posterior nuclear pole (Fig. 4C).

### Inefficient organization of manchette microtubules in *Banf2*^−/−^,*Nup210l*^−/−^ spermatids

Using transmission electron microscopy (TEM) of testis sections, we observed abnormal nuclear development in elongating spermatids in both *Banf2*^+/−^,*Nup210l*^−/−^ and *Banf2*^−/−^,*Nup210l*^−/−^ males when compared to controls. The most striking anomaly in the double mutant was deep invagination of the NE from the caudal pole, visible in approximately 30% of sections (Fig. 5 B, C, D). In cross-sections of uncondensed spermatid nuclei, the invaginated NE appears to be intact and to surround ectopic microtubules (Fig. 5F), but normally positioned manchette microtubules could also be observed parallel to the sectioned NE (Fig. 5H). Nuclear pores were never seen on the invaginated NE. Nuclear pores showed a more restricted localisation at the caudal nuclear pole of uncondensed elongating nuclei in *Banf2*^−/−^,*Nup210l*^−/−^ males, and appeared to be only on one side of the nuclear invagination (Fig. 5G, H) confirming our observations with immunofluorescent localization of NUP153 (Fig 4C). In compacted spermatid nuclei, ectopic microtubules were not evident but nuclear invaginations remained, appearing as large vacuoles (Fig. 5J, K, L).

**Figure 5:**
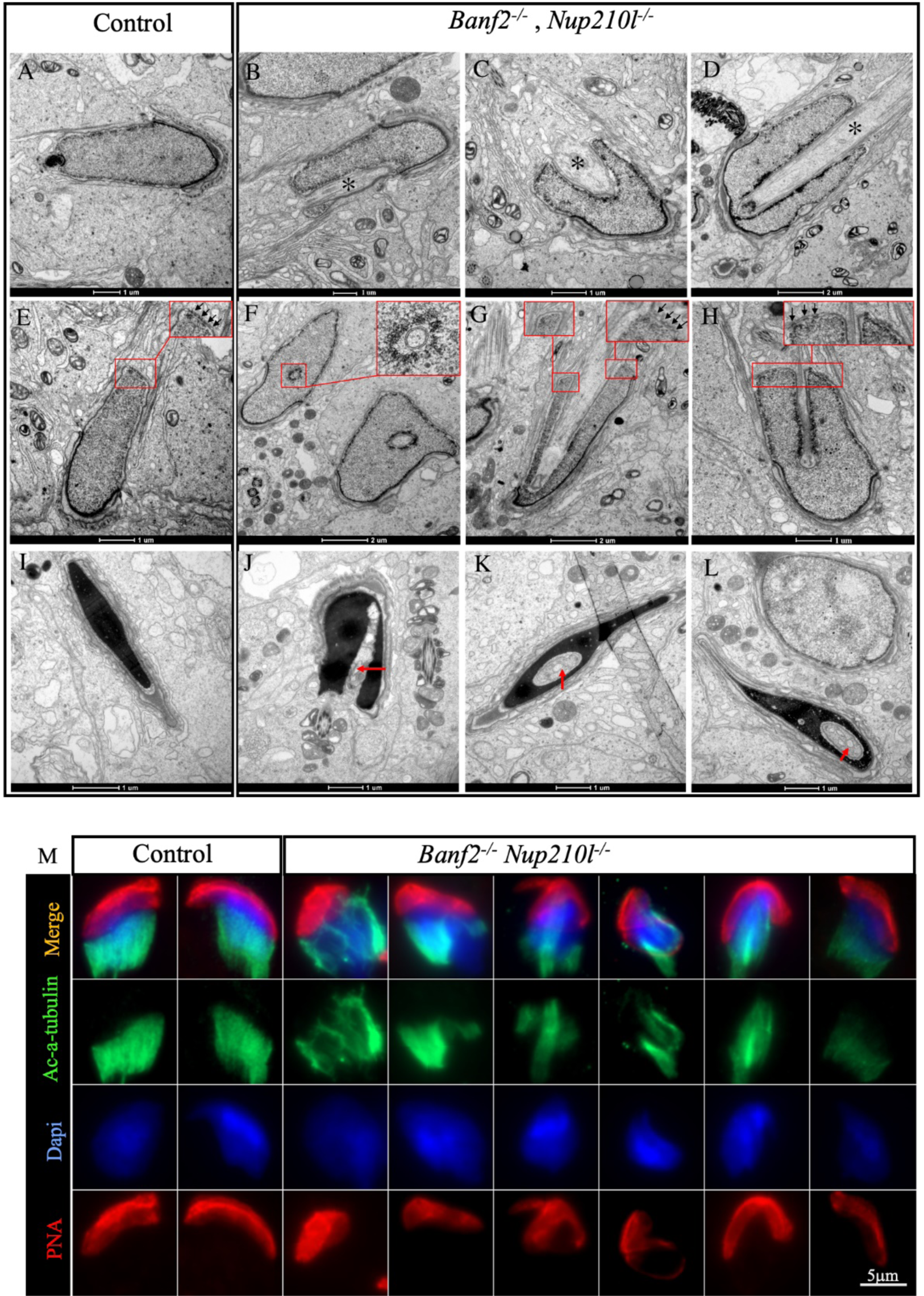
Manchette formation is defective in *Banf2*^−/−^,*Nup210l*^−/−^. Electronic microscopy images of spermatid (step 10–12) from control and mutant mice. As shown by an asterisk (B, C, or D), the *Banf2*^−/−^,*Nup210l*^−/−^ spermatids exhibit nuclear invagination by ectopic microtubules. A transverse section across step 10 spermatid shows that nuclear invagination is composed of microtubular bundles surrounded by a NE (top insert of F). NPC is mostly present on one side of the nuclear invagination, as indicated by the black arrows in the top inserts of (G and H). Step 16 spermatids still have an invagination in the nucleus, the red arrows (J, K, and L) but show that manchette microtubules have depolymerized and been replaced by vacuoles. (M) Immunofluorescence on disaggregated testes using an antibody against acetylated alpha tubulin (Ac-α-tubulin) shows evident disorganization of manchette microtubules.

Further analysis using immunofluorescence with an antibody against acetylated alpha tubulin (Ac-α-tubulin) revealed defective manchette formation in about 45% of *Banf2*^−/−^,*Nup210l*^−/−^ spermatids (Fig. 5M). Defective manchettes, never observed in controls, were characterized by discontinuities in the microtubule sleeve, possibly a consequence of microtubule bundles being diverted from the manchette through a failure to prevent their invagination of the nucleus at its caudal pole.

## Discussion

Our study provides clear evidence that BAF-L and NUP210L are functionally connected genes required for efficient nuclear remodelling in spermatids, and male fertility. Lack of either BAF-L or NUP210L in the mouse is compatible with full fertility (Lu *et al*., 2019; Niu *et al*., 2021; Al Dala Ali *et al*., 2024) although in the case of NUP210L loss there is a drop in spermatozoa quality with half being immotile (Al Dala Ali *et al*., 2024). Here, however, we show that in mice lacking NUP210L, nuclear morphology in round spermatids and manchette formation become sensitive to BAF-L dose, establishing that BAF-L and NUP210L play mutually redundant roles required to maintain the structural integrity of the round spermatid nucleus, to array the NPCs at the caudal pole and to organise microtubules into the manchette. The consequence of insufficient BAF-L in the absence of NUP210L is a catastrophic failure of spermatid nuclear remodelling, leading to male infertility. We reveal function at the caudal NE in spermatids, previously hidden by the functional redundancy of BAF-L and NUP210L.

Although it has been known for over 50 years that NPCs shift to a tight array at the caudal nuclear pole as the manchette forms and spermatid elongation begins (Dooher and Bennett 1973), there has been little notion of how the NPC contributes to spermatid differentiation, beyond controlling traffic into and out of the nucleus. Our study reveals that one of the roles of the NPC array, is to ensure nuclear integrity by regulating the position of polymerising manchette MTs relative to the nucleus. It is not clear how NUP210L and BAF-L attain this, but a similar phenotype of microtubule invagination of the spermatid nucleus from its caudal pole has been reported in rats treated with taxol (Russell *et al*., 1991), and in neurons where the normally axonal tau protein is ectopically positioned close to the nucleus in the cell body (Paonessa *et al*., 2019). Since both taxol and tau stabilise microtubules, the NPC array may normally block stabilisation of MTs that contact the caudal nuclear pole, during the formation of the manchette in step 8 spermatids. This might serve to guide the proto-manchette MTs to the SUN3/SUN4 links in the lateral NE and on to join the sub-acrosomal perinuclear ring (Calvi *et al*., 2015; Gao *et al*., 2020; Pasch *et al*., 2015). Our results show that the spermatid NPC array is an integral part of spermatid NE compartmentalisation, protecting the caudal NE from invagination by MTs as they polymerise to form the manchette. It may achieve this by preventing the diffusion of SUN3/SUN4 MT-tethering links from the lateral NE.

The structural organisation and function of NUP210L has not been studied directly, and can only be inferred from what is known about its paralogue NUP210. NUP210 forms an elastic octamer ring that encircles the nuclear pore membrane where it is proposed to buffer pore function, control pore spacing, and recruit transcription factors to chromatin at the NPC (Zhang *et al*., 2020; Cohen *et al*., 2003; Raices *et al*., 2017). In breast cancer cells, through an interaction with the LINC complex protein SUN2, NUP210 is part of a mechanosensitive connection between the chromatin and the cytoskeleton that is important for metastasis (Amin *et al*., 2021). SUN2 is not present in spermatids, but its close paralogue SUN1, with which it is partially redundant, has been localised to the caudal pole in elongating spermatids along with nesprin-3, a LINC complex partner of SUN1 and SUN2 (Göb *et al*., 2010), and so could link NUP210L to the cytoskeleton. Although mice lacking nesprin-3 are fully fertile, the study of mice lacking nesprin-3 and BAF-L or nesprin-3 and NUP210L should allow this hypothesis to be tested. Thus, in spermatids, NUP210L could have a physical effect on the arrangement of NPCs and their core function, be part of a mechanosensitive chromatin-cytoskeletal link and regulate gene expression. In *Banf2*^−/−^,*Nup210l*^−/−^ round and elongating spermatids, the weakening of nuclear integrity could therefore be a result of the NPC decoupling from both a SUN1-nesprin-3 LINC complex and a chromatin-BAF/BAF-L-cytoskeleton link.

Given that BAF-L can form heterodimers with its paralogue BAF, a conserved chromatin protein, it is most probable that, in the spermatid nucleus, BAF-L is indirectly bound to specific sites on the chromatin through BAF (Tifft *et al*., 2006). The binding of BAF/BAF-L heterodimers (WT fertile mice) or BAF homodimers (*Banf2*-KO fertile mice) to DNA may enable the resulting chromatin to link to the cytoskeleton through the NPC. Nevertheless, how BAF or BAF-L connect to the NPC or the NE in spermatids is not clear. Most of the proteins known to link BAF to the NE in somatic cells were not detected in human or rodent spermatids (lamin A/C, emerin, LEMD3, LEMD2), or do not localise to the caudal pole (Vester *et al*., 1993; Elkhatib *et al*., 2017; Elkhatib *et al*., 2015). A candidate protein that might interact with BAF-L, is the spermatid-predominant protein, LEMD1, that has an N-terminal LEM-domain and a C-terminal transmembrane domain. In human, LEMD1 labels the nucleoplasm in round spermatids, shifting to the caudal pole as the spermatid elongates (Elkhatib *et al*., 2017). As a possible explanation for the observed functional redundancy of NUP210L and BAF-L in maintaining spermatid nuclear integrity, we propose that the NPC normally participates in two distinct chromatin-cytoskeleton links: the first involving NUP210L, SUN1-nesprin-3 and possibly BAF, and the second a nucleoporin other than NUP210L and BAF-L.

In human, the infertile man lacking NUP210L has severe oligozoospermia and produced immotile spermatozoa (100%) with large uncondensed heads (83%) and retained histones (100%) (Arafah *et al*., 2021). The *Banf2*^−/−^,*Nup210l*^−/−^ and the *Banf2*^+/−^,*Nup210l*^−/−^ mice presented here do not reproduce the human sperm phenotype exactly but there are similarities that may indicate common consequences of compromised NPC function during spermiogenesis: severe but incomplete spermatogenic blockage during nuclear remodelling, epididymal spermatozoa with uncondensed heads (14%), histones retained in spermatozoa (25%) and few motile sperm (1-2%). Based on our findings in the mouse, the severity of the human *NUP210L*-KO phenotype indicates that a biological process able to complement the loss of NUP210L function is not active in the patient. This could be the result of a recent genetic mutation (digenism) or an ancient mutation common to all humans. One candidate for the former, carried by the patient, is a rare potentially pathogenic missense variant (p.Pro485Leu) in NUP153, a nucleoporin present at the spermatid caudal nuclear pole (Al Dala Ali *et al*., 2024).

Our double knockout model reveals an example of functional redundancy in a previously unknown biological process indispensable for spermiogenesis. Genetic redundancy has most commonly been evidenced between paralogues, but rarely between unrelated genes, since it is practically impossible to identify genes involved in distinct redundant pathways in the absence of sequence homology. Nevertheless, there are many reports of conserved testis predominant genes whose inactivation in the mouse does not appear to have any effect on fertility in the laboratory environment, indicating that genetic redundancy is common during spermatogenesis (Khan *et al*., 2018; Miyata *et al*., 2016; Lu *et al*., 2019) As is clearly illustrated by our findings, functional redundancy renders biological processes invisible to molecular genetics based on single-gene inactivation. Furthermore a large study of almost 1000 azoospermic men only identified plausible monogenic causes in 20% of their cohort (Nagirnaja *et al*., 2022). Digenism should therefore be considered when designating candidate causal genetic variants in the context of human male fertility. Firstly, even in an infertile man with an obvious gene inactivation, mutation of an unrelated second gene may underlie the phenotype, and this could confound attempts to validate the variant as causal in the mouse. Secondly when filtering variants from exome sequencing data under a digenic model the allelic frequencies that predict less than 1:10 000 affected individuals are high and would be excluded from monogenic screens: *e.g.,* 1:10 for autosomal recessive mutations in each gene and 1:100 for autosomal dominant or X-linked mutations. Thirdly, in human cases the pathogenic combination of gene variants is already created, limiting possibilities. Identifying the causal variants will of course be arduous at such high allelic frequency cut-offs, requiring mouse models, but should be investigated in best case scenarios involving unambiguous LoF mutations in testis-specific genes. The potential rewards are to uncover biological processes that presently cannot even be imagined.

## Material and Methods

### Generation of knockout mice

Knockout alleles in *Banf2* and *Nup210l* were created by non-homologous end-joining repair of a Cas9 generated double strand break as previously described for *Nup210l* (Al Dala Ali *et al*., 2024). The specific guide RNA sequence (+PAM) used to target *Banf2* was CCCATGTCTGTTGCAGAAGA (+TGG). We selected the allele *Banf2^em42Mmjm^* with a deletion of the first 23 nucleotides from exon 3 - AAGATGGACGACATGTCGCCCAG. The alleles were created in the C57BL/6NCrl strain. Double mutants were created from mice produced by backcrossing to the outbred Swiss strain. The mice presented in this study are on a mixed C57BL/6NCrl-Swiss background. The necessary approval for our study was obtained from our regional animal experimentation ethical committee, CEEA14.

### RT-PCR analysis of testis transcripts from the *Banf2*^em42Mmjm^ allele

Transcripts were amplified with primers in exon 1: o5845f - ACCCGCCGTTACAGATCCCAG and exon 4: o5847r – TCAGGCAGGTGGAGCTCTGG. After agarose electrophoresis, the amplified bands were cut from the gel, purified and sequenced with the same primers.

### Sperm analysis

For sperm analysis, the cauda epididymis of each mouse was dissected into small pieces using two scalpels to allow motile sperm to swim out. These pieces were then incubated in 2 ml of medium (DMEM, HEPES - 22320022 Thermofisher) at 37°C for 10 minutes. Sperm suspensions were obtained, and the motility of the sperm was assessed by direct microscope observation. The total number of sperm was estimated using a Malassez hemocytometer, and the morphology of the sperm was evaluated after staining with SpermBlue stain (Microptic, Barcelona, Spain), following the standard protocol. The morphology of the head and tail was assessed individually for each mouse, with a minimum count of 100 cells per sample.

### Testis and epididymis histology

Mice testes and epididymis were obtained from both control and mutant mice. They were fixed in Bouin’s solution at room temperature overnight. The fixed tissues were then dehydrated, embedded in paraffin, and sectioned to a thickness of 7 µm for the testis and 5 µm for the epididymis using an Ultrathin Microtome from Leica, Germany. The testis sections were stained with H-PAS staining (Sigma-Aldrich), while the epididymis sections were stained with Mayer’s hematoxylin and eosin from Sigma-Aldrich, following a standard protocol

### Immunofluorescence - Testis sections

Testis samples were collected and fixed in 4% paraformaldehyde (PAF) at 4 °C overnight. The fixed testes were dehydrated in a graded sucrose solution, embedded in OCT, and cut into 5 μm sections. The sections were then rehydrated with phosphate-buffered saline (PBS) for 5 minutes, permeabilized with 0.3% Triton X-100 in PBS for 30 minutes, and blocked with 1% BSA, 7% NGS, and 0.1% Triton X-100 for 1 hour at room temperature. The sections were incubated overnight with primary antibodies in the blocking solution. After three washes in a 0.1% Triton X-100 solution, the slides were incubated with secondary antibodies in the blocking solution for 1 hour at room temperature. Following 3 washes with 0.1% Triton X-100 in PBS, the nuclei were counterstained with 100 ng/ml DAPI in PBS, and the slides were mounted using fluoromount-G mounting media (Southern Biotech).

### Immunofluorescence - Testis cell spreads

To isolate individual cells, the testes were retrieved, and the tunica albuginea removed. The testicular pulp was dilacerated using two scalpels. The cells obtained were suspended in 7 ml of DMEM, HEPES medium (Thermofisher). They were then washed twice with PBS, fixed in a 2% paraformaldehyde solution for 5 minutes, rinsed twice again with PBS, and finally spread onto slides using a Cytospin (Shandon). The slides were permeabilized with 0.5% Triton X-100 in PBS for 15 minutes and blocked with a solution of 1% BSA and 7% NGS at room temperature for 1 hour. Immunofluorescence was then performed as described above for testicular sections.

### Antibodies

The following primary antibodies were used for immunofluorescence investigations at the specified dilution: anti-Lamin B1 (ab151735 - Abcam) 1:100, anti-acetylated histone H4 06866 - Upstate) 1:400, and anti-NUP153 (ab24700 - Abcam) 1:50, anti-Ac-α-tubulin (T7451 - Sigma) 1:20, and anti-BAF (ab129184 - Abcam) 1:100.

Acrosomes were labeled directly with Lectin PNA Alexa Fluor™Plus 594 (L32459 - Thermofisher) 1:600. Secondary antibodies were from ThermoFisher: Goat anti-Rabbit IgG Alexa Fluor™Plus 488 (A32731) 1:400; Goat anti-Mouse IgG Alexa Fluor™Plus 488 (A11001) 1:400. Donkey anti-Rabbit IgG Alexa Fluor™ Plus 647 (Invitrogen-A32795) IF-1:400

### Terminal deoxynucleotidyl transferase dUTP nick-end labeling (TUNEL) assay on testes

The testes were collected and fixed with 4% PAF at 4°C overnight, dehydrated, and embedded in paraffin. They were then sectioned to a thickness of 5 μm using a microtome. The resulting slides were heated at 55°C for 30 minutes, followed by deparaffinization and rehydration. Subsequently, the slides were incubated with a freshly prepared solution containing 0.1% Triton and 0.1% sodium citrate for 20 minutes at 37°C. After two rinses with PBS, the slides were incubated with a blocking solution (3% BSA, 10% NGS in 10mM Tris-HCl, pH 7.6) for 45 minutes at room temperature. Following this, the slides were incubated with a Tunnel mixture (*In Situ* cell detection kit, Roche) for 1 hour at 37°C. After three washes with PBS, the nuclei were stained with a solution of 100 ng/ml DAPI in PBS and mounted using Fluoromount-G mounting media (Southern Biotech).

### Chromomycin A3 staining

The sperm was washed twice with PBS, then fixed in a mixture of methanol and acetic acid (3:1 v/v) for 30 minutes at 4°C. The sperm was spread onto slides and allowed to air-dry. Each slide was stained by applying a solution of chromomycin A3 (0.25 mg/ml) in McIlvain buffer at pH 7.0 for 20 minutes. Afterward, the slides were rinsed twice in McIlvain buffer. The nuclei of the sperm were counterstained with a solution of Hoechst (0.5 g/ml) for 5 minutes and rinsed twice with PBS. Fluoromount-G mounting media was used for mounting the slides. Slides were examined using a fluorescence microscope (ApoTome2, Germany).

### Transmission electronic microscopy

The testes were fixed overnight at 4°C using a solution of 2.5% glutaraldehyde and 2% PFA in 0.1M sodium cacodylate buffer. The following day, the testes were washed three times in sodium cacodylate buffer for 10 minutes each and then post-fixed in 2% osmium tetroxide in 0.1M cacodylate buffer for 1 hour at room temperature. After the post-fixation, the testes were washed three times in buffer for 15 minutes each and dehydrated in a series of ethanol concentrations (25%, 50%, 75%, and 100%). Subsequently, the testes were embedded in epoxy resins. Ultrathin sections of 70 nm thickness were obtained using a Leica UC7 instrument and deposited on slot grids. These sections were contrasted with 1% aqueous uranyl acetate for 10 minutes and lead citrate for 4 minutes. The resulting grids were observed using an FEI Tecnai G2 microscope operating at 200 KeV, and the acquisition of images was performed using a Veleta camera from Olympus, Japan.

### Statistical Analysis

The significance of the variance between the means in study groups was assessed using Student’s t-test. A *p*-value below 0.05 was considered significant.

## Acknowledgements

We thank Nicolas Brouilly and Aïsha Aouane for help with electron microscopy analyses performed at PiCSL-FBI core facilty (IBDM, AMU-Marseille), member of the France-BioImaging national research infrastructure. We are very grateful to Stéphane Zaffran, Gaëlle Odelin and Adeline Spiga-Ghata of the MMG Animal House Facility for enabling our mouse work. We also thank Keltouma Ali Nehari and Paul Picot for attentive animal care.

## Data availability

All relevant data can be found within the article and its supplementary information.

