## Supplementary Figures S1-S5 for "Nucleoporin NUP210L and BAF-paralogue BAF-L together ensure microtubule organization and nuclear integrity in spermatids"

**Supplemental material**

Supplemental material consists of five Figures:

Supplemental_Fig_S1: Creation and validation of the *Banf2*$^{em42Mmjm}$ null allele (*Banf2*$^{-}$).

Supplemental_Fig_S2: Normal spermatogenesis and fertility in *Banf2*$^{-/-}$ knockout males.

Supplemental_Fig_S3: PAS-H stained testicular sections showing disruptions in spermatogenesis in the *Banf2*$^{-/-}$,*Nup210l*$^{-/-}$ mice, characterized by the presence of vacuoles and a decrease in the number of elongated spermatids.

Supplemental_Fig_S4: The nuclear envelope is not deformed in round spermatids isolated from the seminiferous tubules of *Banf2*$^{-/-}$,*Nup210l*$^{-/-}$ males.

Supplemental_Fig_S5: Localization of BAF in epididymal sperm of control and *Banf2*$^{-/-}$,*Nup210l*$^{-/-}$ mice.

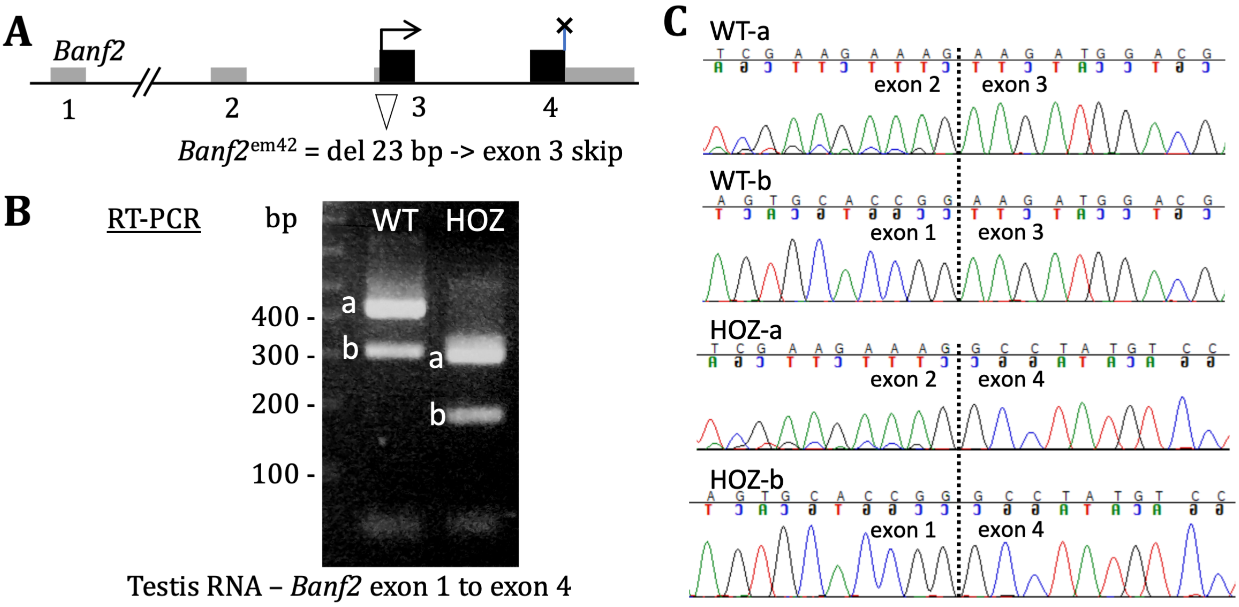


**Figure S1:** Creation and validation of the *Banf2*$^{em42Mmjm}$ null allele (*Banf2*$^{-}$). (A) Diagram of the mouse *Banf2* gene locus showing the deletion of the first 23 bp of exon 3 in the *Banf2*$^{em42Mmjm}$ allele. Coding exonic sequence is in black boxes, non-coding in grey. Exons and introns are not drawn to scale. (B) RT-PCR of adult testis RNA from a WT male and a male homozygous for *Banf2*$^{em42Mmjm}$ (HOZ). *Banf2* normally produces two transcripts, one with all exons (a) and one without the non-coding exon 2 (b). Both WT and HOZ produce two bands but the bands are shorter in HOZ than in WT. (C) The two bands (a and b) amplified from WT or HOZ testis RNA were each purified and sequenced. The chromatograms for the sequence of the distinct exon junctions are shown. This shows that the *Banf2*$^{em42Mmjm}$ allele produces transcripts with and without exon 2, and that both these transcript types lack exon 3.

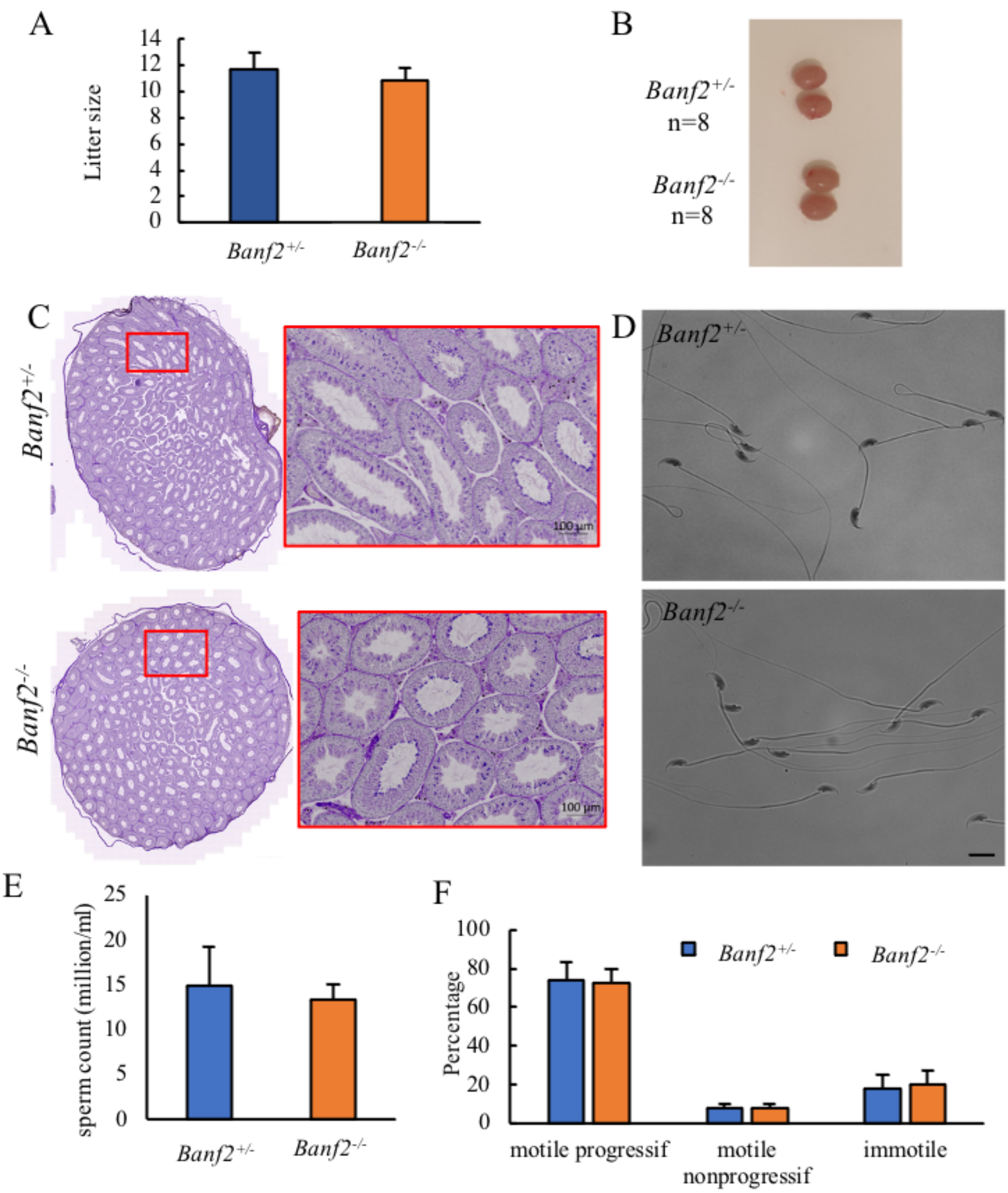


**Figure.S2:** Spermatogenesis is normal in *Banf2*$^{em42Mmjm/em42Mmjm}$ knockout mice. (A) Litter size produced by *Banf2* $^{-/-}$ mice is comparable to that of *Banf2* $^{+/-}$ males (70 d*pp*). *p*-value = 0.29. Average number of litters per male is 4.5 for *Banf2* $^{+/-}$ and 4.75 for *Banf2* $^{-/-}$, *p*-value = 0.54. Four males per genotype were mated with two females each for 12 weeks. (B) Testes from 3-month-old WT (above) and *Banf2*$^{-/-}$ males (below) demonstrate the normal testicular size in *Banf2*$^{-/-}$ mice. (C) Testis histology stained with periodic acid-Schiff (PAS) reveals normal organization and development of germ cells in the seminiferous tubules of *Banf2* mice. (D) Epididymal sperm stained with a SpermBlue show normal morphology in *Banf2*$^{-/-}$ mice. Scale bar 20μm (E) There is no observed reduction in sperm count in *Banf2*$^{-/-}$ mice. N = 6. *p*-value = 0.1. (F) Quantitative analysis of the mobility of *Banf2*$^{-/-}$ epididymal sperm indicates no significant difference compared to WT littermates. *p*-value = 0.7 for motile sperm, 1 for motile non-progressive sperm, and 0.59 for immotile sperm. N = 6.

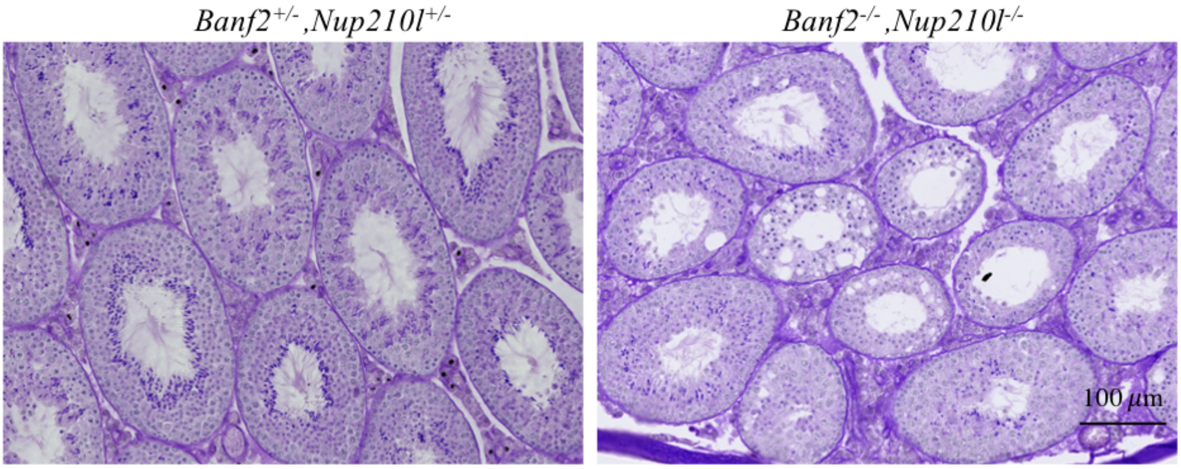


**Figure S3**: PAS-stained testicular sections revealed disruptions in spermatogenesis in the *Banf2*$^{-/-}$,*Nup210l*$^{-/-}$ mice, characterized by the presence of vacuoles and a decrease in the number of elongated spermatids.

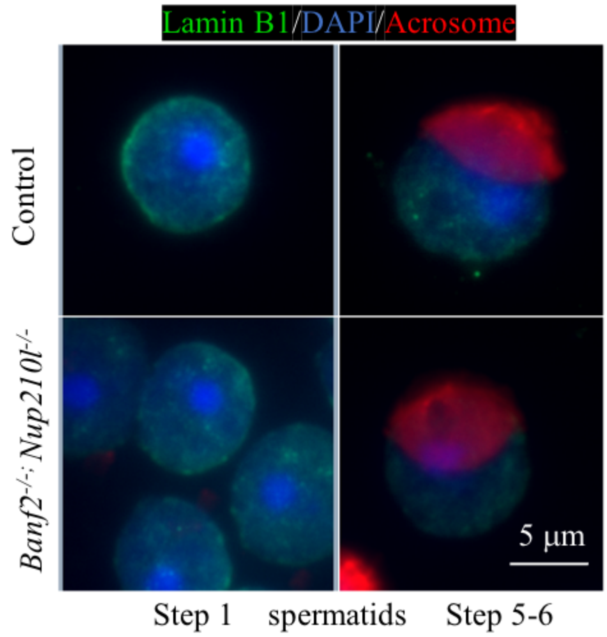


**Figure S4**: The nuclear envelope is not deformed in round spermatids isolated from the seminiferous tubules. The nuclear envelope of round spermatids at step 1 and step 5-6 is shown labelled with an antibody against Lamin B1, in control and double mutant males. The nuclear DNA is labelled with DAPI and the spermatids are staged by labelling the acrosome with fluorochrome-conjugated lectin. Spermatids were harvested from adult testis that were dilacerated to liberate cells from the seminiferous tubules and generate a suspension of germ cells.

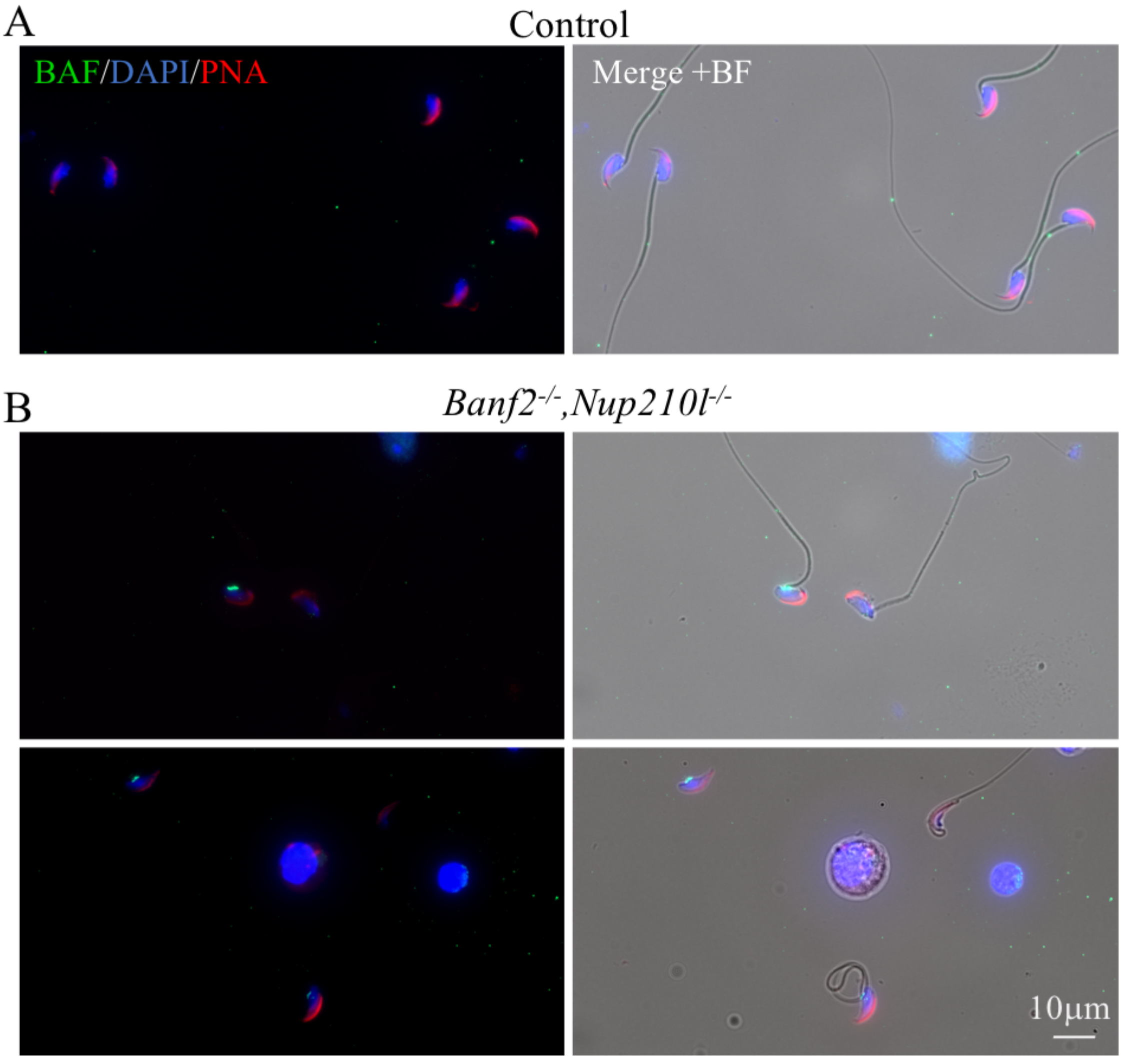


Figure S5: Localization of BAF in epididymal sperm of control and *Banf2*$^{-/-}$,*Nup210l*$^{-/-}$ mice. (A) Complete absence of BAF signal (green) in control spermatozoa. (B) retention of BAF signal at the posterior nuclear pole in 23% of mutant spermatozoa (n=100). The nucleus is counterstained with DAPI (blue), and the acrosome is stained with PNA (red).
